# The conserved regulatory basis of mRNA contributions to the early *Drosophila* embryo differs between the maternal and zygotic genomes

**DOI:** 10.1101/769638

**Authors:** Charles S. Omura, Susan E. Lott

## Abstract

The gene products that drive early development are critical for setting up developmental trajectories in all animals. The earliest stages of development are fueled by maternally provided mRNAs until the zygote can take over transcription of its own genome. In early development, both maternally deposited and zygotically transcribed gene products have been well characterized in model systems. Previously, we demonstrated that across the genus *Drosophila*, maternal and zygotic mRNAs are largely conserved but also showed a surprising amount of change across species, with more differences evolving at the zygotic stage than the maternal stage. In this study, we use comparative methods to elucidate the regulatory mechanisms underlying maternal deposition and zygotic transcription across species. Through motif analysis, we discovered considerable conservation of regulatory mechanisms associated with maternal transcription, as compared to zygotic transcription. We also found that the regulatory mechanisms active in the two genomes, maternal versus zygotic, are quite different. For maternally deposited genes, we uncovered many signals that are consistent with transcriptional regulation through control at the level of chromatin through factors enriched in the ovary, rather than precisely controlled gene-specific factors. For genes expressed only by the zygotic genome, we found evidence for previously identified regulators such as Zelda and GAGA-factor, with multiple analyses pointing toward gene-specific regulation. The observed mechanisms of regulation are consistent with what is known about regulation in these two genomes: during oogenesis, the maternal genome is optimized to quickly produce a large volume of transcripts to provide to the oocyte; after zygotic genome activation, mechanisms are employed to activate transcription of specific genes in a spatiotemporally precise manner. Thus the genetic architecture of the maternal and zygotic genomes and the specific requirements for the transcripts present at each stage of embryogenesis determine the regulatory mechanisms responsible for transcripts present at these stages.

**Author summary:** Early development in animals is a unique period of time, as it is controlled by gene products from two different genomes: that of the mother and that of the zygote. The earliest stages of development are directed by maternal mRNAs and proteins that are deposited into the egg, and only later does the zygote take over the transcription of its own genome. In this paper, we use data from 11 fruit fly species characterizing all the genes transcribed by the mother and later by the zygote, to investigate how transcription is regulated in the maternal and zygotic genomes. While we find some conserved regulatory elements at both stages, regulation of maternal transcription is much more highly conserved across species. We present evidence that maternal transcription is controlled in large co-regulated chromatin domains, while zygotic transcription is much more gene-specific. These results make sense in the context of where these genes are being transcribed, as maternal transcripts are generated in support cells which churn out a large amount of mRNA during oogenesis, while zygotic genes are often transcribed in a particular time and place in the embryo.

## Introduction

Development is a sequential process, where each step builds on the one before it. The earliest stages of embryonic development are therefore critical, as processes such as cleavage cycles and the beginnings of axial patterning become the basis for all subsequent developmental processes. Regulation of these important tasks is controlled by mRNAs and proteins, and perhaps unsurprisingly then, mRNA levels in *Drosophila* are found to be precisely controlled during early embryogenesis[1, 2]. This precise control of transcript levels is especially remarkable, given that the transcripts at early stages of development come from two different genomes. The first set of transcripts are those deposited into the egg by the mother, while the second set are transcribed from the zygotic genome[3–5]. However, the regulatory mechanisms responsible for this precise control are not yet fully understood.

During oogenesis, the oocyte itself is mostly transcriptionally silent[6]. Instead, support cells called nurse cells synthesize RNA, proteins, and organelles which are transported into the oocyte[7]. These maternally produced mRNAs are responsible for many of the critical events of early embryogenesis, such as the rapid cleavage cycles, the establishment of body axis, and the coordination of the handoff of control to the zygotic genome. This handoff of developmental control from mother to zygote, known as the maternal to zygotic transition (MZT), is complex from a regulatory standpoint. Critical housekeeping genes retain a steady transcript level, despite changing the genome of origin. New transcripts must be synthesized from the newly activated zygotic genome, and maternal transcripts must be degraded, in a highly regulated and time-specific manner[8]. This transition is well studied in model systems such as *Drosophila melanogaster*, where maternal mRNA degradation regulators such as *smaug* (*smg*)[8] and regulators critical to the activation of the zygotic genome such as *zelda* (*zld*)[9, 10] have been identified. When the transition of developmental control between the two genomes is complete, the zygotic genome must be poised to carry out the rest of development in a precise manner. One process that exemplifies the precision required at the handoff to the zygotic genome is segmentation in *Drosophila*. This process begins with broad maternal gradients which control transcription of early zygotic gap genes, and later pair-rule genes, at precise locations within the embryo at specific developmental times[11, 12].

Regulation of transcripts in development has been the subject of considerable study in *D.melanogaster*. Much of this study has been focused around the process of the MZT or other important events in early development, such as patterning along the anterior-posterior or dorsal-ventral axes. For example, a number of regulators of maternal transcript degradation at or prior to the MZT have been identified[13–16]. Zygotic transcription activation has also been the subject of considerable study, and has implicated critical transcription factors such as *zelda* and *grainy head*[4]. How transcripts are transported into eggs has been the subject of some study[7,17,18], as has how those maternal transcripts are regulated post-transcriptionally[3,19–23]. Post-transcriptional mRNA regulation is especially crucial at the maternal stage as new transcripts cannot be produced after the completion of oogenesis. However, how transcript production is regulated in the nurse cells is largely unknown. As transcript pools at both the maternal and zygotic stages are highly conserved over evolutionary time[24], we employed a comparative approach to investigate gene regulation at these stages.

In this study, we uncover regulatory elements that are associated with transcription in the early *Drosophila* embryo, from both maternally deposited and zygotically transcribed genes. We use motif analysis to compare regulation of maternal versus zygotic transcription, and also investigate how regulation at these two stages is different across *Drosophila* species. To this end, we used a previously generated RNAseq dataset from Atallah and Lott, 2018, which sampled embryos from a developmental stage where all transcripts are maternal (stage 2[25, 26]) and a stage after zygotic genome activation (end of stage 5 [23, 24]), across 14 species, representing ∼50 million years of divergence time. Here, we used the transcript abundance data from 11 of these species (due to limitations in genome annotation quality, see Methods), representing the same span in divergence time, to examine putative regulatory regions of maternally deposited or zygotically transcribed genes. Through comparisons of these sequences and associated gene transcription levels, we identified a number of sequence motifs as being enriched in either maternally deposited or early zygotically expressed genes. We found a high similarity between motifs across all species, suggesting a high level of conservation for regulation of transcription within each genome (maternal and zygotic). At the stage controlled by maternal transcripts, we found a high number of motifs that bind to proteins annotated with insulator function or that have previously been associated with boundaries between topologically associating domains (TADs). Our findings suggest that maternal transcription is largely controlled through regulation of chromatin state, and not through gene-specific mechanisms. Many transcription factors predicted to bind the identified motifs were found to be enriched in ovaries[27]. After zygotic genome activation (stage 5), we find many of the motifs known to be associated with early zygotic transcription, such as the binding site for the pioneer transcription factor, Zelda, reinforcing many previously identified aspects of transcriptional regulation at this stage. We also find a larger number of motifs with less significant enrichment at this stage, with evidence that points to these motifs regulating a smaller subset of genes. This study provides evidence for global control of maternal transcription at the level of chromatin, while zygotic transcription is regulated in a more gene-specific manner. This is especially striking considering that the maternal transcript pool is more highly conserved than that of the early zygote[24].

## Results

### Discovered maternal-associated motifs are bind architectural proteins; discovered zygotic-associated motifs bind to known zygotic regulators

To examine the regulatory basis of maternal and zygotic transcription, we surveyed the genomes of 11 *Drosophila* species for regulatory elements. These species represent the evolutionary divergence of the *Drosophila* genus, encompassing divergence times from 250,000 to 50 million years[28]. The RNAseq datasets produced from Atallah and Lott (2018)[24] were used. These data sampled two developmental stages, one where all transcripts present are maternally derived (stage 2, Bownes’ stages[25, 26]) and the other after zygotic genome activation (the end of stage 5, or the end of blastoderm stage). The transcript abundance data was used to classify each gene as being on or off at both stage 2 and stage 5 for each species (see Methods). For each gene, we extracted sequences at likely locations for proximal regulatory elements (see Methods). To accommodate the varying annotation quality of the various species, this search encompassed introns, exons, and a 2kb region upstream of the gene.

To identify motifs associated with maternally deposited genes, we employed HOMER[29]. For most species, a characteristic pattern emerged where the most enriched motifs were present in the upstream region of the maternally deposited genes, with less enriched motifs appearing in exons (Fig S1). Some motifs, possibly representing repressor binding sites, were enriched in the upstream and intron region of genes that were not maternally deposited as compared to the genes that were maternally deposited (Fig S1).

Analyzing regulatory elements at the post-zygotic genome activation stage (stage 5) presents a challenge, as it is difficult to distinguish newly transcribed zygotic mRNAs from residual maternally deposited mRNAs. At this stage, roughly half of the transcripts present are maternal transcripts that have not yet been degraded[8,30–32]. Therefore, to interrogate regulatory elements associated with zygotic transcription, we restricted our search to genes that do not have transcripts present at stage 2 but do have transcripts present by stage 5. Because of these stricter requirements for zygotically transcribed genes, there were far fewer genes in the dataset (66,206 genes in the stage 2 dataset combined from all species, compared to 10,215 total genes in the stage 5 dataset for all species), resulting in a reduction in statistical power. However, without these assumptions, we risk failing to identify signals associated specifically with zygotic transcription amongst the signal of maternal transcription.

To determine which proteins are likely to bind to maternal or zygotic motifs, we used Tomtom[33] to evaluate the similarity of the discovered motifs to several motif databases (Table 1) for *D. melanogaster*. The motifs found in maternally deposited transcripts are similar to those discovered previously in two different contexts: those associated with topologically associated domains (TADs)[34], and those associated with housekeeping promoters[35, 36]. This is consistent with existing data showing that functions of maternally deposited genes are enriched for genes with housekeeping activities[36, 37]. In order to determine whether the motifs associated with maternal transcripts in our data were simply due to the inclusion of promoter elements from housekeeping genes, we measured the enrichment of these motifs in maternally deposited genes that are not housekeeping genes (see Methods). We found that our motifs are strongly enriched (p < 1e-34) in maternally deposited genes even when excluding housekeeping genes (S6 Figure A). This indicates that these motifs are having a strong effect outside that of those contained in housekeeping genes during this stage. Thus, we hypothesize that the regulatory mechanisms responsible for generating TADs[34] are also responsible for maternal transcripts, and that maternal transcription may be regulated by the establishment of TADs. TADs are genomic regions where the chromatin on one side of the boundary interacts substantially less than expected with the chromatin on the other side, and interactions of DNA elements within the domains can be promoted. While TADs are generally thought to be associated with transcription [34], there is some controversy as to the nature and magnitude of the effect of TADs on gene expression [38], as disruption of TADs has not been found to be sufficient to alter transcription in some cases.

**Table 1:**
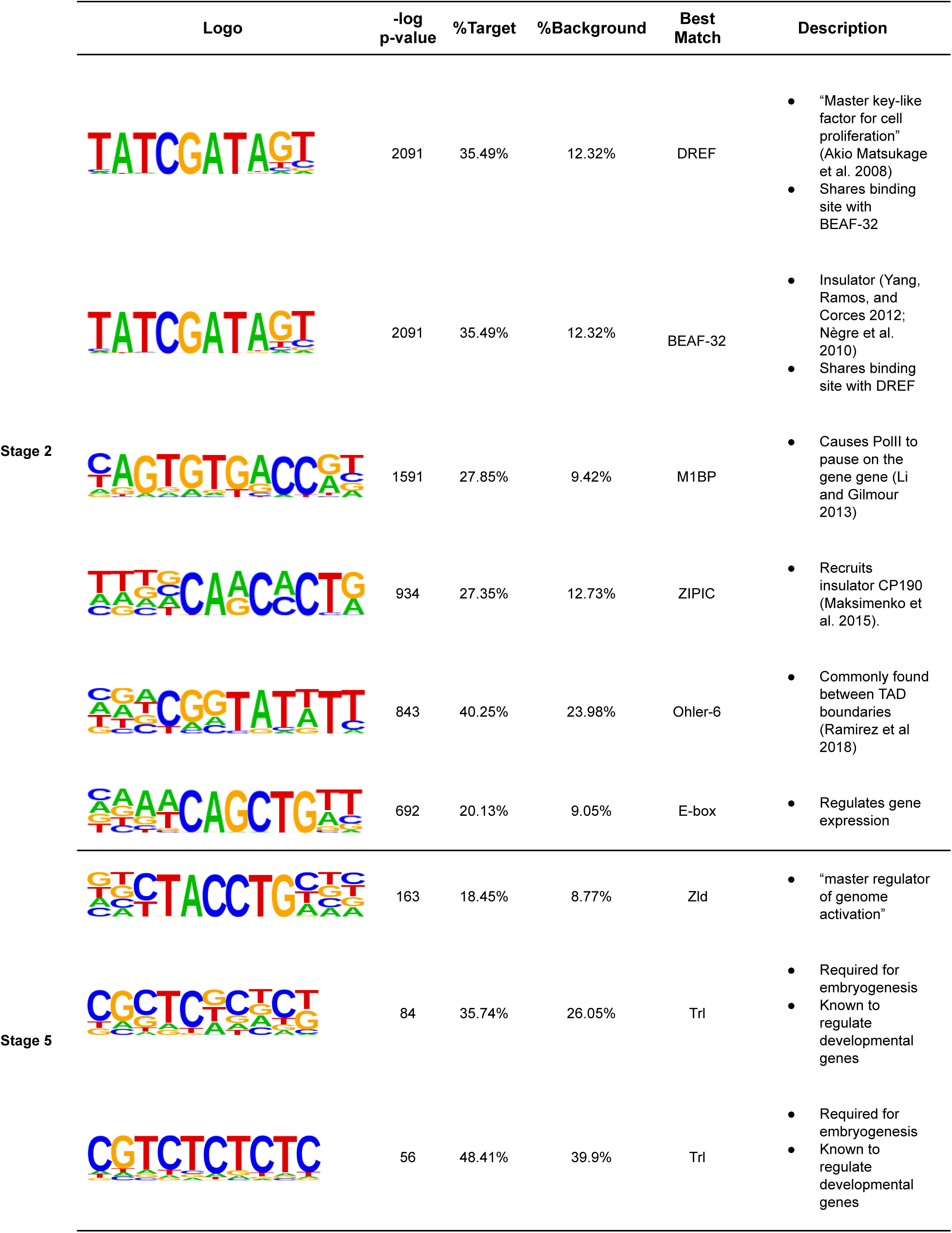
A summary of the top ranked motifs. HOMER was used to find motifs enriched in the 2kb windows upstream of maternally deposited genes (stage 2) and zygotically transcribed genes (stage 5). Sequence logo shows the consensus motif where the probability of each base is proportional to its representative character. P-value is given by HOMER. %target represents the percent of either maternally deposited or zygotically expressed genes that contain at least one instance of the motif. %background indicates the percent of all genes that contain this motif. Best match indicates protein with a previously identified binding site that mostly closely matches the discovered motif (see Methods).

The motifs associated with maternally deposited genes are predicted to bind several insulators or architectural proteins. An insulator is a regulatory element that suppresses the interactions of other regulatory elements with genes, or prevents the spread of chromatin state. An architectural protein is a protein that organizes and regulates chromatin structure. The most prominent motif by q-value binds to DNA replication-related element factor (DREF), a known architectural protein and the “master key-like factor for cell proliferation”[39]. It is required for normal progression through the cell cycle. It is known to occur in the promoters of many cell proliferation genes and to interact with chromatin remodeling proteins. Interestingly, DREF binding site overlaps with the binding site for BEAF-32, another well-researched protein that acts as an insulator[40, 41] that often appears between head-to-head genes (genes with adjacent promoters that get transcribed in opposite directions). Another identified motif is predicted to bind ZIPIC, which is known to bind and recruit CP190, an insulator. A previous study provides evidence for the co-localization of ZIPC and BEAF-32[42], which likely work together with CP190 to perform insulator functions. Thus of the most enriched motifs in maternal genes (DREF, BEAF-32, ZIPC), many have previously identified roles as insulators or in other ways regulating chromatin state.

Another maternal motif identified is predicted to bind M1BP (motif-1 binding protein), which causes RNA polymerase II (Pol II) to pause on the gene[43]. Pol II pausing is critical to early zygotic expression[36, 44] but its function in producing the maternal transcriptome is unknown. Several functions have been suggested for this Pol II pausing behavior, including maximizing transcription speed once certain conditions are met, synchronizing with RNA processing machinery, reacting to other developmental or environmental signals, keeping chromatin accessible, and acting as an insulator. Given that M1BP is both maternally deposited at high levels and has increased expression in the early embryo, it is possible that M1BP has multiple functions at different time points. During oogenesis, pausing to wait for external signals or RNA processing machinery seems counterproductive to maximizing transcription in the ovary, but the other function of maintaining a state of open chromatin and solidifying TAD boundaries may be very important. In contrast, at stage 5 it may be much more important to maximize expression in response to certain signals.

In searching for motifs associated with zygotic expression, we recovered motifs for well-known regulators of the zygotic genome (Table 1). We only identified a small number of highly enriched motifs at this stage, and thus were able to predict a much smaller number of predicted factors binding to these motifs, including Trl (or GAGA factor) and Zelda. Trl is a known early zygotic activator and chromatin remodeler[45–47] and Zelda is known as a “master key regulator” to early developmental genes[9, 48] and appears to be a pioneer transcription factor that establishes the initial chromatin landscape of the zygotic genome[5]. In addition to these high-quality motifs, we found a large number of motifs with lower quality scores (Table S1). These motifs may regulate spatio-temporal specific genes that we observe in the early embryo, and thus have a lower enrichment score due to our whole-embryo approach being ill-equipped to finding such specific patterns.

### Similar motifs appear in different species

To quantify the conservation of the discovered motifs across the 11 species in our study, we used Tomtom[33] to measure the similarity between the sets of motifs discovered in different species. For a motif to be considered conserved between two species, we required that it be discovered by HOMER in both species and for Tomtom to report a statistically significant alignment score (see Methods). At the maternal stage, we found that high quality (q-value < 1e-100 by HOMER, see Methods) motifs tended to be well-conserved (Fig 1A) with a large percentage of the total discovered motif content shared across species. We observed that sister species *D. pseudoobscura* and *D. persimilis* are unique in that they have the highest number of motifs that are either species-specific or are only shared with each other, and have the fewest number of motifs shared with the rest of the species. This is especially noteworthy considering that this lineage is roughly in the middle of the distribution of divergence times from most of the other species, and thus many more distantly related species comparisons have a higher degree of motif conservation than do any comparisons with these two species. This is consistent with previous results[24] that this lineage has a disproportionately high number of changes in transcript abundance for its phylogenetic position, and suggests that these large number of changes in transcript abundance may be due to the large scale changes in regulation in these species observed here. When comparing the rest of the species, we found a relatively higher number of conserved motifs shared between pairs of species within the *Drosophila melanogaster* species group (*D. melanogaster*, *D. simulans*, *D. sechellia*, *D. yakuba*, *D. erecta*, *D. ananassae*), and a slightly reduced number of conserved motifs between the *D. melanogaster* group species and the more distantly related species (*D. willistoni*, *D. mojavensis*, *D. virilis*) (Fig 1B). At stage 5, we do not observe a high percentage of conserved motifs between species, rather we observe many motifs that are significantly enriched in just one or two species. We also observe little phylogenetic signal in the data, with the only detectable pattern being that the species with the longest divergence time from the rest of the species, *D. virilis* and *D.mojavensis*, have slightly fewer shared motifs (Fig 1 C,D). If the unique motifs at either stage indeed represent newly evolved regulatory mechanisms, we expect that these motifs to be rare or to have a smaller frequency difference between transcribed and non-transcribed genes. Either of these effects would raise the false discovery rate as reported by HOMER, which makes the number of species-specific zygotic motifs identified all the more remarkable. Additionally, more highly conserved motifs should require less power to be discovered as they are by definition present across more species, and thus we should have more power to identify them than less-conserved motifs. It is still possible that there are more conserved motifs at the zygotic stage that we do not observe due to the lower number of genes used at this stage. Despite this, however, the dominant signal we find from the motifs we have power to detect is non-conserved. This is underscored by the observation that when we reduce our quality threshold for motifs at stage 5, we still do not observe motifs to generally be conserved across species (Fig S4 B).

**Fig 1:**
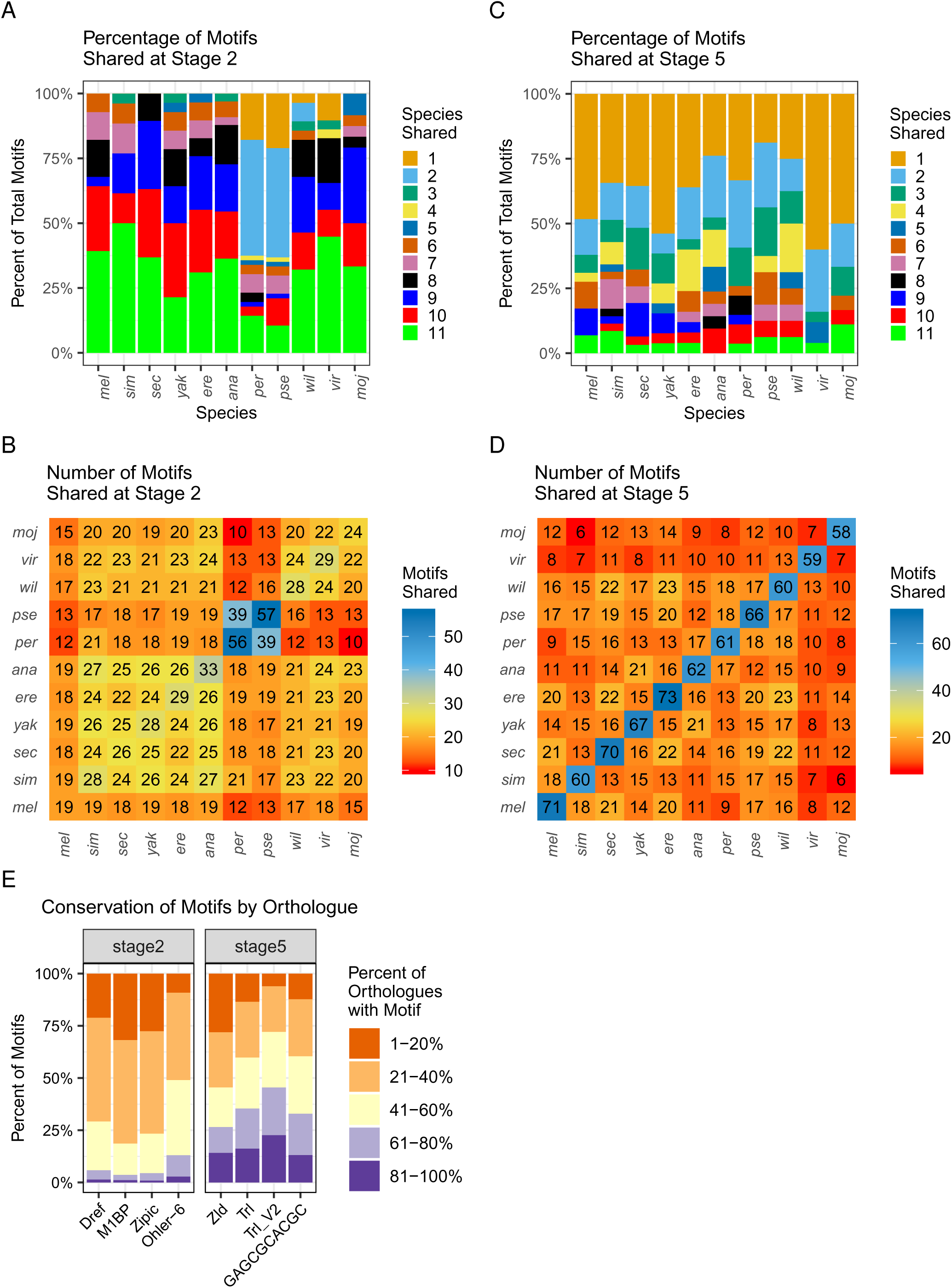
Motifs associated with maternal deposition are largely shared across species, zygotic motifs are likely to be species-specific. For each analysis represented in A-D, motif enrichment was determined for each group of genes at each stage (all maternally deposited genes at stage 2; or zygotic genes at stage 5) separately in each species, then lists of enriched motifs at each stage were compared across species. For stage 2 motifs, we required motifs to have a −log qvalue > 100, while for stage 5 motifs we required motifs to have a −log qvalue > 10 (see Methods). (A, C) Percent of motif content in the upstream region that is found to be shared between species at stage 2 and stage 5, respectively. The number of species that share each motif is indicated by the color of the bar. Note that in stage 2, a large majority of motifs are shared in all (11 species) or almost all (9 or 10 species), with the exception of *D. pseudoobscura* and *D. persimilis*, sister species that share common motifs between themselves but are different from the rest of the species. Zygotic motifs identified at stage 5 are much more likely to be species specific or shared by only a couple of species. (B, D) Number of motifs shared between each pair of species at stage 2 and stage 5, respectively. Comparisons of one species to itself indicate the total number of motifs that fit quality criteria discovered in that species. Comparing the number of shared motifs between pairs of species, there is some signal of the phylogeny in stage 2 (B), with *D. melanogaster* subgroup species sharing more motifs in common with one another than they do with the more distantly related species, and *D. pseudoobscura* and *D. persimilis* with the highest number of motifs in common but the most differences from the remaining species. For stage 5 (D), apparent patterns include both the number of species-specific motifs (diagonal) and less apparent phylogenetic structure. (E) Conservation of top motifs in orthologous genes across species. Y-axis indicates all of the instances of the motif of interest within the upstream region. Coloration represents how many species’ orthologues also contain that motif. In general, top motifs at the zygotic stage (stage 5) are more likely to be conserved in orthologous genes at this stage. This sets up a contrast with parts A-D, where maternal deposition is broadly associated with a shared set of motifs across species, but part E shows that orthologous maternal genes are less likely to share a specific motif.

### Motif conservation by gene

While these results show that some motifs are important to regulation in the genomes of multiple species, do not speak to whether orthologous genes in different species tend to contain similar motifs. To investigate whether regulation was conserved at the level of individual genes, we compared the motif content of each *D. melanogaster* gene (see Methods) to the motif content of each of its orthologs from other species. We counted motifs as conserved between two species if the motif appeared in both orthologs. For both stage 2 and stage 5, we categorized motifs based on the percent of orthologs for which the motif was conserved (Fig 1E). Motifs have different levels of gene-specific conservation between stages, with maternal stage motifs appearing to have lower conservation across orthologues than zygotic stage motifs, where a larger proportion of orthologues possess the same motif. This is striking, as this seems to imply that while gene expression and regulation are both highly conserved for maternal genes, which genes are regulated by a particular regulator is not. It is possible that the genes that are missing motifs compared to their orthologues are regulated by different motifs, or that the same motifs that are in radically different positions in different species. As many different maternal motifs appear to be regulating transcription at the level of chromatin state, these motifs may be able to function interchangeably. Thus this environment may be more conducive to more motif turnover at this stage but with higher conservation of transcription overall[24], as compared to the zygotic stage.

### Motif position

While similar binding motifs identified in multiple species implies that regulatory proteins with similar binding domains are acting in these species, we can also verify the similarity in the regulatory machinery by the relative positions of the binding sites relative to the genes they are regulating. To investigate whether the discovered motifs had the same positional relationship with the transcription start site (TSS) across all species, we generated position frequency data for each motif. For each gene, we examined each position starting from 2kb upstream of the TSS to the 3’ end of the gene body, and whether there was a motif at that position. Many of the most prominent motifs shared a similar distribution pattern, characterized by a strong peak at −100bp, and sometimes a secondary peak at −340bp (Fig S2). To quantify this similarity, we performed an Anderson-Darling test on each motif for each pair of species, which indicated that 65% (stage 2) and 91% (stage 5) of motif distributions are identical between species (percent of motifs for which p < .05). This suggests conservation of the relationship between binding to these motifs and initiation of transcription. The higher conservation of motif position in stage 5, which has fewer conserved motifs between species than stage 2, may be consistent with this stage having more gene-specific regulation, as discussed further below.

### Motif Strandedness

While some studies focus on finding motifs with a particular orientation relative to their proximal genes[49], there is some evidence that motifs do not behave in a strand-specific manner[50]. To evaluate the importance of the strandness of the discovered motifs, we generated a regression to predict expression level that differentiated between forward and reverse versions of each motif (see Methods). This regression indicated a significant difference between the forward and reverse versions of many motifs. For example, we found the E-box motif affects the log-odds of maternal deposition by .192 in the forward orientation but only .115 in the reverse orientation (t-test, p < .001). For almost all motifs, different strands had the same qualitative effect on expression, but with different magnitudes, indicating that while motifs had the same effect regardless of orientation, their efficiency could be increased if the orientation was optimal.

While the strandedness of motifs may play a small role in their overall effect, we want to know if strandedness makes a qualitative difference to our motifs effects on transcript level, and if we can use motif strand to improve our model. To determine this, we ran HOMER exclusively on the same strand that the gene appeared on, rather than the default mode of scanning both strands. This resulted in the same set of motifs being discovered. This is consistent with the regression results that show that each motif, whether located on the positive strand or the negative strand relative to the transcription start site, has the same qualitative effect on gene expression, indicating that the direction of each motif had minimal effect on expression. To evaluate whether the strand the motif was located on relative to the gene was predictive in whether a gene was transcribed at a particular stage, we constructed another regression using only the data from the same-strand motifs. This regression performed less well than the regression using motifs from both strands (AIC = 7915.8 for the unstranded regression, AIC = 8612.5 for the stranded regression for a representative species *D. ananassae*). Overall, this suggests that motif binding elements need not bind in a strand specific manner to induce their effects, though the optimal orientation provides measurable increase in their effect on transcription. This result is the same at both stage 2 and stage 5.

### GO analysis

While we have identified a set of motifs that together seem to be responsible for early embryonic RNA content, we next asked if these motifs are likely to be regulating genes with specific types of functions. To this end, we performed gene ontology (GO) analysis on groups of genes, based on their motif content. To simplify this analysis, we chose to focus on the top 8 motifs as reported by HOMER, and for each of the 8 motifs, we performed GO analysis on the transcript pools at each stage as well as on each motif individually[51, 52]. We initially performed a GO analysis on both the maternally deposited and zygotically transcribed transcript pools, disregarding motif content. When comparing stages, we observe no overlap between GO terms (Fig 2A), which is consistent with our expectations that the genes that are activated in the zygote have different functionality to those transcripts that are maternally deposited, especially as our definition of zygotically transcribed genes excludes genes present in stage 2. When examining genes containing specific motifs within each stage, we observe that many of the stage 2 motifs show a similar pattern in the GO categories they are associated with, with the strongest associations belonging to the DREF motif, which is strongly associated with most identified categories (Fig 2B). This could be an indication that there is a high degree of homogeneity in terms of the types of genes these motifs may regulate. In contrast, the stage 5 motifs present in zygotic-only genes show more variety in the GO terms of genes they are associated with (Fig 2C), which could be indicative of more specific regulation for these genes at this stage.

**Fig 2:**
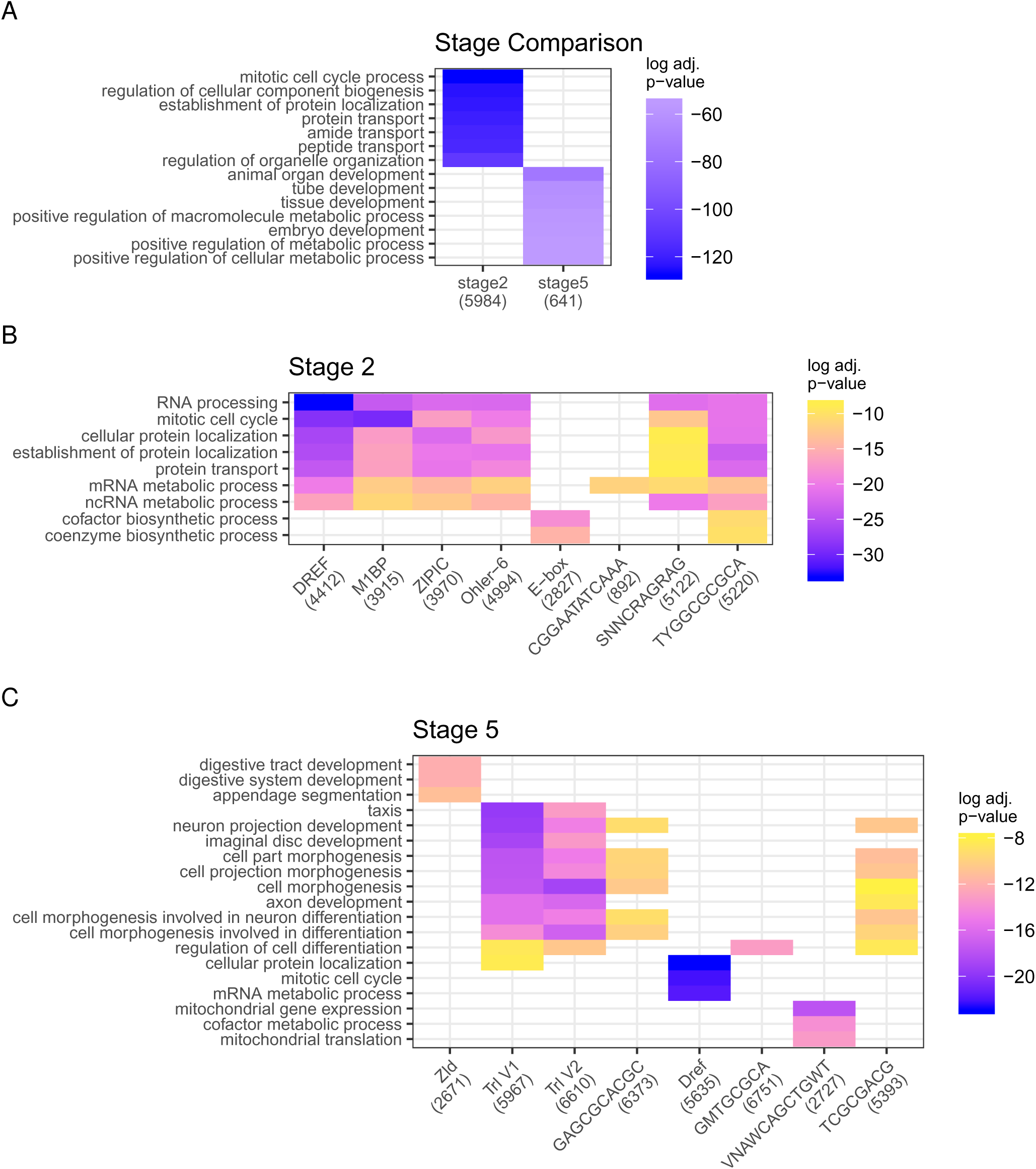
Top GO terms show that motifs regulate broader set of genes at the maternal stage, and a more specific set of developmentally associated genes at the zygotic stage. (A) GO terms associated with each stage. Note that the set of identified GO categories does not overlap between stages. (B) GO terms associated with top motifs in stage 2, where a majority of motifs are associated with similar broad GO categories (C) GO terms associated with top motifs in stage 5, some motifs are associated with the same categories, some appear to be more specialized, with identified categories showing more specificity than categories associated with stage 2

While the previous GO analysis indicated that the top motifs at stage 2 display significant overlap in associated GO categories, this does not exclude the possibility that specific GO categories are regulated by specific motifs. To search for more specific motifs, we performed motif analysis using HOMER to find overrepresented sequences in the top GO terms within maternally deposited genes, resulting in several motifs which are enriched in specific GO terms (Fig. S5), though very few of them are significantly enriched after multiple test correction. These motifs do not appear in other analyses, and do not have strong matches to proteins expressed in the ovary found in the literature. Because these motifs are associated with a small subset of genes, we hypothesized that these motifs confer specificity to transcription of specific genes with accessible chromatin. To determine whether these motifs are associated with increased expression at stage 2, we used linear models to measure the effect of the presence of these motifs, specifically in genes that already contain motifs that bind to architectural proteins, or whose adjacent genes are highly expressed. We did not find that the presence of these GO term-specific motifs increased the odds of maternal deposition (Fig S5). It is possible that this result is due to the lack of statistical power surrounding these motifs, as these motifs are somewhat rare. This result could also be the underlying biology, however, and these motifs could be non-functional at stage 2.

### Predicted maternal motif binding proteins are enriched in the ovary

Next, we investigated whether the potential motif binding proteins we identified were plausible regulators of maternal deposition. It is unclear whether the motifs we identified as enriched in maternally deposited genes are associated specifically with maternal deposition, given that chromatin regulators are important at all stages in all tissues. To investigate, we used modENCODE[53] transcript abundance data to compare the mRNA transcript levels for proteins predicted to bind our discovered motifs, and found increased expression in ovaries (Fig 3A) as compared to other tissues sampled. This pattern exists, though to a lesser extent, in the FlyAtlas 2 dataset[54], which is a tissue-specific database of transcript levels that utilizes RNA-seq data rather than microarray analysis. The discrepancy between the two datasets could be due to the differences in gene expression measurement method or in experimental methods. The transcripts for these proteins also show moderately high abundance in our own dataset (File S5). While it has been demonstrated that mRNA levels do not necessarily mirror protein levels[55], the enrichment of mRNA in ovaries compared to other tissues is reasonable evidence that these proteins are important in ovaries.

**Fig 3:**
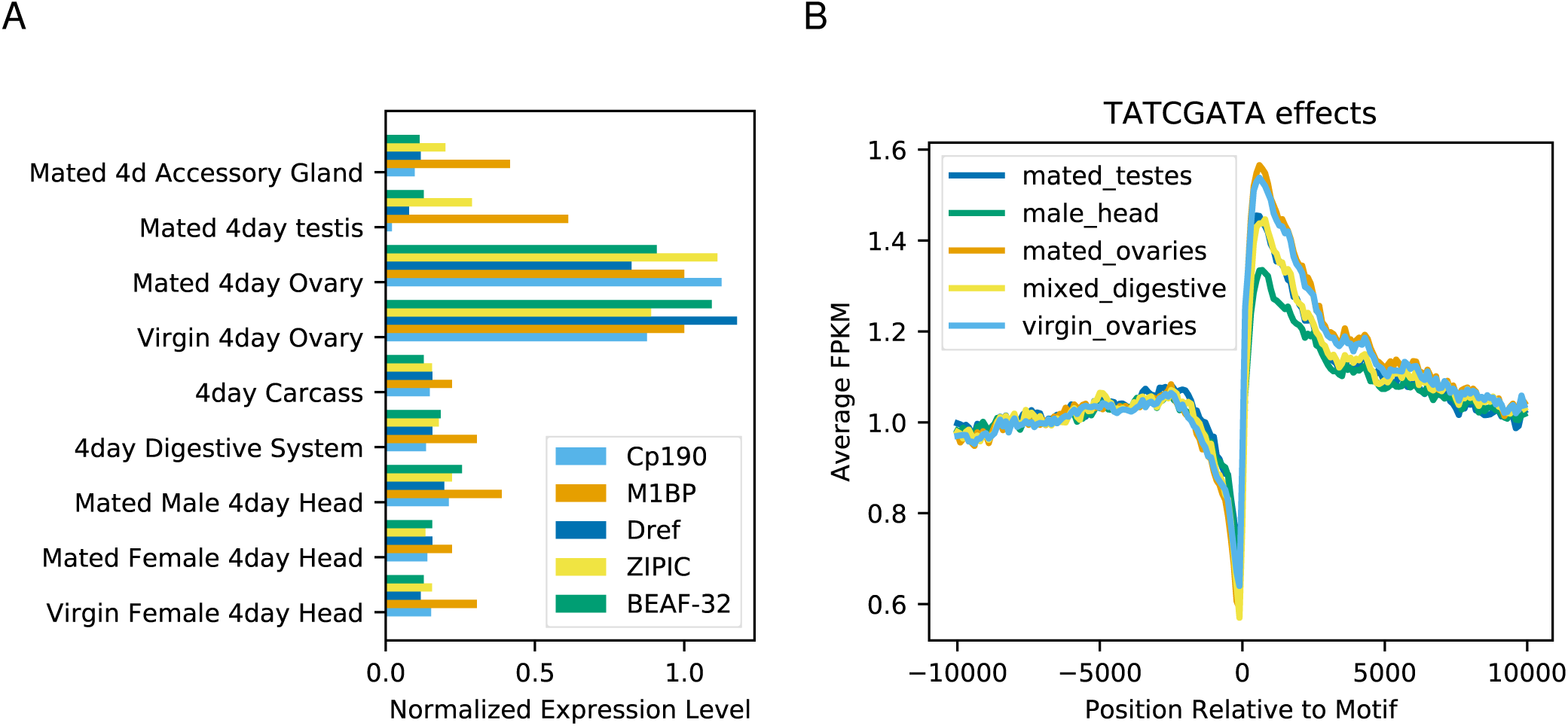
Identified maternal regulators are ovary enriched, as is their effect on transcription (A) RNA levels of putative binding proteins by tissue type. Transcript abundances within each gene have been normalized such that the average abundance in ovaries is equal to 1. While identified maternal regulators have regulatory functions in multiple tissue types, they are highly enriched in ovaries compared to other tissues. (B) Average normalized expression levels versus proximity to motif by tissue type. Normalization was performed by dividing each expression value by the average expression from 9.9-10kb away. While binding sites for identified maternal regulators are present in multiple tissues, the effect on gene expression is stronger in ovaries compared to other tissues.

To investigate whether these proteins are acting to affect transcription in the ovaries specifically, we examined the expression profiles of RNA in various tissue types (referenced in Fig 3B) from existing RNA quantification datasets[53, 56]. For each instance of a motif of interest, we extracted the transcript level from within a 20kb window surrounding the motif and measured the normalized relative transcript level for each position (an example of this is shown in Fig 3B). While the relative normalized transcript level changes in each of the measured tissues, the effect is strongest in ovaries, indicating that the presence of one of these binding sites is associated with a higher increase in transcript levels in the ovary compared to other tissues.

As the motifs associated with maternal transcription also act to some degree in other tissues, we next wanted to ask whether the motifs were more enriched in maternally deposited genes than in genes expressed in other tissues. To determine whether regulation in different tissue types were associated with different motifs, we ran HOMER in the same manner as with the maternal stage data to discover enriched motifs (see Methods) in transcripts present in other tissues, as identified from ModENCODE data[57]. We found that most other tissue types were also enriched in the same motifs discovered in transcripts present in stage 2 embryos. However, examining the frequency of motifs in specific genes revealed that the majority of those motifs were from genes that were shared between those tissue types and stage 2 embryos. When we exclude genes that are expressed in stage 2 embryos, HOMER fails to identify the original set of motifs as enriched in male larval gonads, male reproductive tract, adult heads and adult midgut. Furthermore, HOMER detects the motifs at a lesser rate in larval ovaries, larval CNS, and intestinal tract. Despite being identified in fewer tissue types and at a lesser rate in other tissue types as compared to the stage 2 expression levels, the observation that these motifs may also have important functions in other tissue types is consistent with the literature. For example, DREF is known to be important for cell proliferation and chromatin regulation, and is active in many other tissues[58, 59]. These motifs are likely associated with many housekeeping genes that are vital to a variety of tissue types.

### Maternally deposited genes are physically clustered on the genome

In addition to motifs, we observed several other effects that were related to early embryonic RNA content. Given that many of our discovered motifs bind architectural proteins, we hypothesize many effects may be linked to the physical location of genes on the chromosome. We examined the positional distribution of transcribed genes in various tissue types (Fig 4A). As previous papers utilizing the Hi-C method have shown correlation with active topologically associated domains (TADs) and gene expression[60, 61], we predicted that any tissue type where regulation is dominated by architectural proteins to transcribe a set of genes physically clustered on the chromosome. To compare the physical gene clustering of transcription at the maternal stage with that of other tissue types, we acquired several RNAseq datasets from NCBI/GEO[62] and performed a Wald–Wolfowitz runs test[63] on each tissue of the previously described tissue types. While all tissues examined showed a strong preference for groupings of transcribed genes, embryonic stage 2 samples were the most highly grouped (Fig 4B). This result was robust to changes in the threshold of what is considered to be expressed (see Methods). This pattern of physical co-expressed gene clustering on the chromosome is consistent with our model of regulation via architectural proteins.

**Fig 4:**
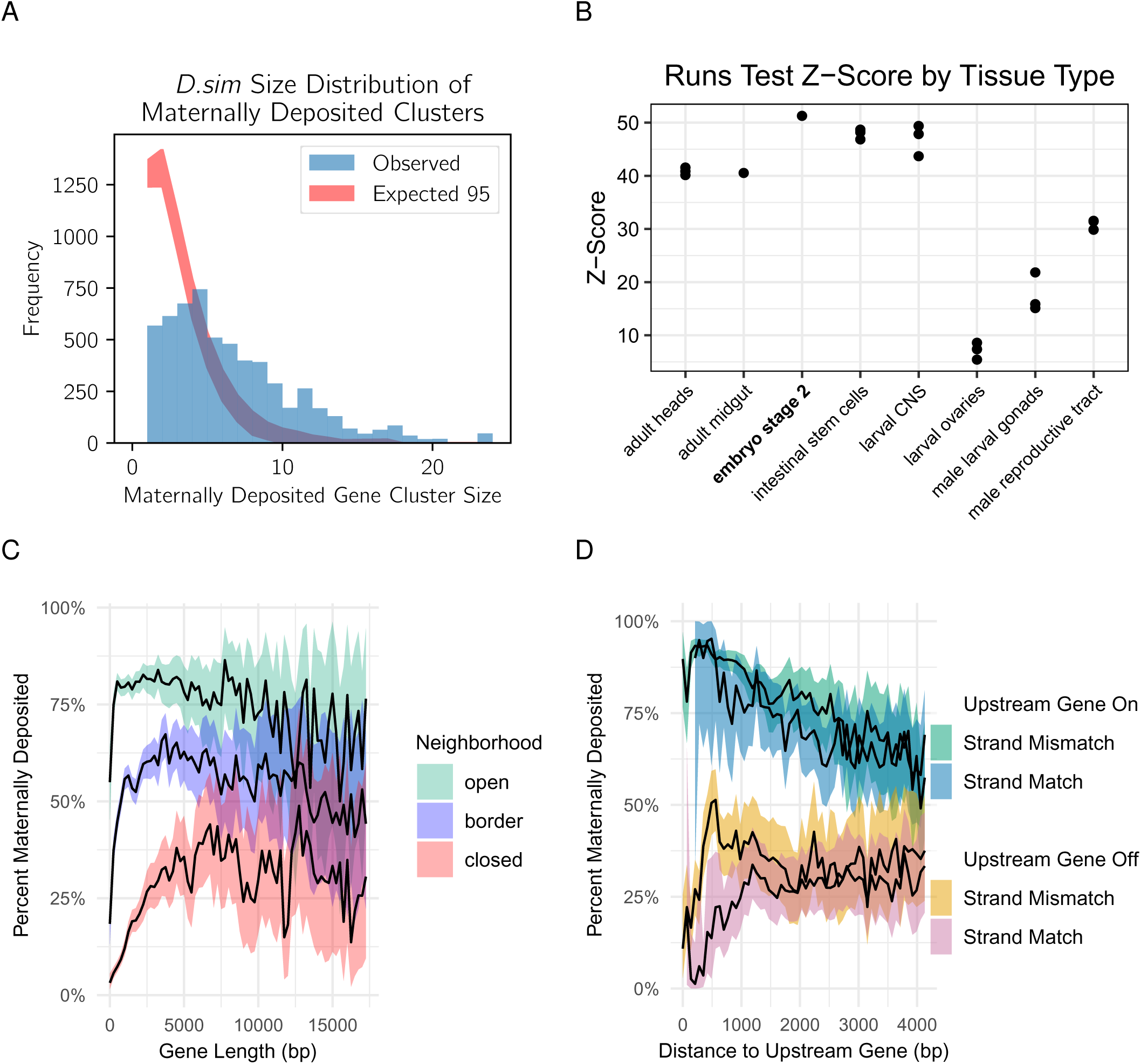
Chromatin stage and maternal deposition. For the analyses in A and B, genes were categorized as either expressed or not expressed (see Methods) and adjacent expressed genes were considered to be clustered, with a cluster size equal to the number of constituent genes. (A) Physical clustering of maternally deposited genes along the chromosome, in a representative species (*D. simulans*). The shaded blue region represents the observed frequency of co-expressed maternal gene clusters of various sizes. The red region represents the 95% CI constructed with 10,000 bootstrap iterations. Maternal genes are co-expressed in clusters along the chromosome more often than expected, given the percent of the genome that is transcribed at this stage. (B) Physical clustering of co-expressed genes on chromosomes in various tissue types. In order to compensate for differing proportions of the genome that are expressed in each tissue type, physical clustering was measured by performing a Wald-Wolfowitz runs test and taking the z-score (see Methods). Maternally expressed genes, represented by stage 2 embryos, show the highest proportion of physical clustering of co-expressed genes, though other tissues such as intestinal stem cells and larval CNS also have highly physically clustered co-expressed genes. (C) Gene length by number of adjacent maternally expressed genes, “open” indicating both adjacent genes are expressed, “border” indicating that one is expressed, and “closed” indicating that neither are expressed. Genes that with more expressed neighbors are more likely to be maternally deposited, regardless of length. Genes without expressed neighbors are less likely to be maternally deposited, with the odds increasing as length increases. (D) Odds of maternal deposition versus distance to the nearest upstream gene by upstream expression and strand. Distance is measured by from transcription start site (TSS) to TSS. When the upstream gene is maternally deposited, odds of maternal deposition are high, but decrease with distance regardless of strand. When the upstream gene is not maternally deposited, odds of maternal deposition are low and have a strand-dependent relationship with distance.

While these results speak to the pattern of clustering of expression for maternal genes in terms of adjacent genes being on or off, they do not account for the distance between genes. To answer the question of whether this clustering phenomenon is dependent on distance, we examined the distance to adjacent genes. We observed a trend whereby proximity to an active promoter increases the odds of maternal deposition (Fig 4C). This effect was slightly affected by the strandedness of the two genes whereby genes that have an opposite orientation are more likely to have different expression. This is consistent with observations from previous studies[34] that consecutive genes on the same strand were more likely to show co-expression, while consecutive genes on opposite strands were more likely to have different expression.

Many previous studies have observed that zygotic genes tend to be short in length[24,30,64,65]. In addition to affecting transcription speed, shorter gene lengths result in a smaller distance between transcriptional units along the chromosome, especially when considering which strand the gene is on. To explore gene length in maternal genes and the relationship between gene length and the position on the chromosome, we measured the maternal deposition rates with respect to gene length. We observed a trend that in most species, shorter genes are less likely to be maternally deposited. There are differences in the length of maternal genes across species, and this trend could be partly due to the bias for more highly annotated genomes to be enriched in shorter genes (Fig 4D). Additionally, chromatin context seems to heavily influence this effect: when the adjacent genes are off, gene length is much more important (Fig 4C) and very short genes are very likely to be off. This could be because shorter genes are more likely to be influenced by the regulatory machinery of a nearby gene. Alternatively, longer genes might be long enough to physically isolate themselves more effectively and establish their own unique regulatory environment.

Given that a number of motifs found in this study are bound by proteins annotated as insulators, and the motifs are similar to those that are associated with TADs, we asked where the motifs found in our dataset can be found relative to TAD boundaries. Previous results suggest that architectural proteins are prevalent in the centers of TADs as well as the boundaries[34], and may be involved in mediating interactions of the DNA within a TAD [38]. To determine the location of motifs in the context of TADs, we assessed the transcription of nearby genes relative to the transcription of a gene with these identified motifs. For each regulatory region, the gene nearest to that regulatory region was examined, as well as two genes downstream and two upstream. The frequency of motifs was measured based on the transcript abundance pattern of these five genes. Many of the top motifs including Dref, M1BP, Zipic, and E-box, occur more frequently in the center of maternally deposited gene clusters, rather than on the edge of clusters. (t-test p-values 7e-3, 2e-6,3e-10, and 1e-4 respectively). This is consistent with previous results[34], and may suggest an important role for architectural proteins in promoting interactions within a TAD as well as potentially in establishing TAD boundaries.

### Stage-specific genes are isolated on the genome

Given that maternally deposited genes are physically clustered together in the genome, we wanted to examine if this pattern held with the set of genes that were stage-specific. To determine if consecutively expressed cluster size is related to stage-specificity of transcript representation, we examined maternal-only (transcripts present at stage 2 and entirely degraded by stage 5) and zygotic-only genes (transcripts present at stage 5, not present at stage 2; for both stage-specific categories, see Methods for further definitions) and their frequencies in clusters of different sizes. We determined that for most species, in contrast to all maternally deposited genes, both maternal-only and zygotic-only genes are more likely to be in smaller (1-3 consecutive active genes) groups than in larger groups (more than 3 consecutive active genes) (Fig 5, A and B). For these stage-specific genes, this could be an indication that control of stage-restricted genes is more specific, affecting single genes rather than larger clusters. Results for most other analyses of maternal-only genes were unable to be obtained due to the very low number of genes in this category (see Methods).

**Fig 5:**
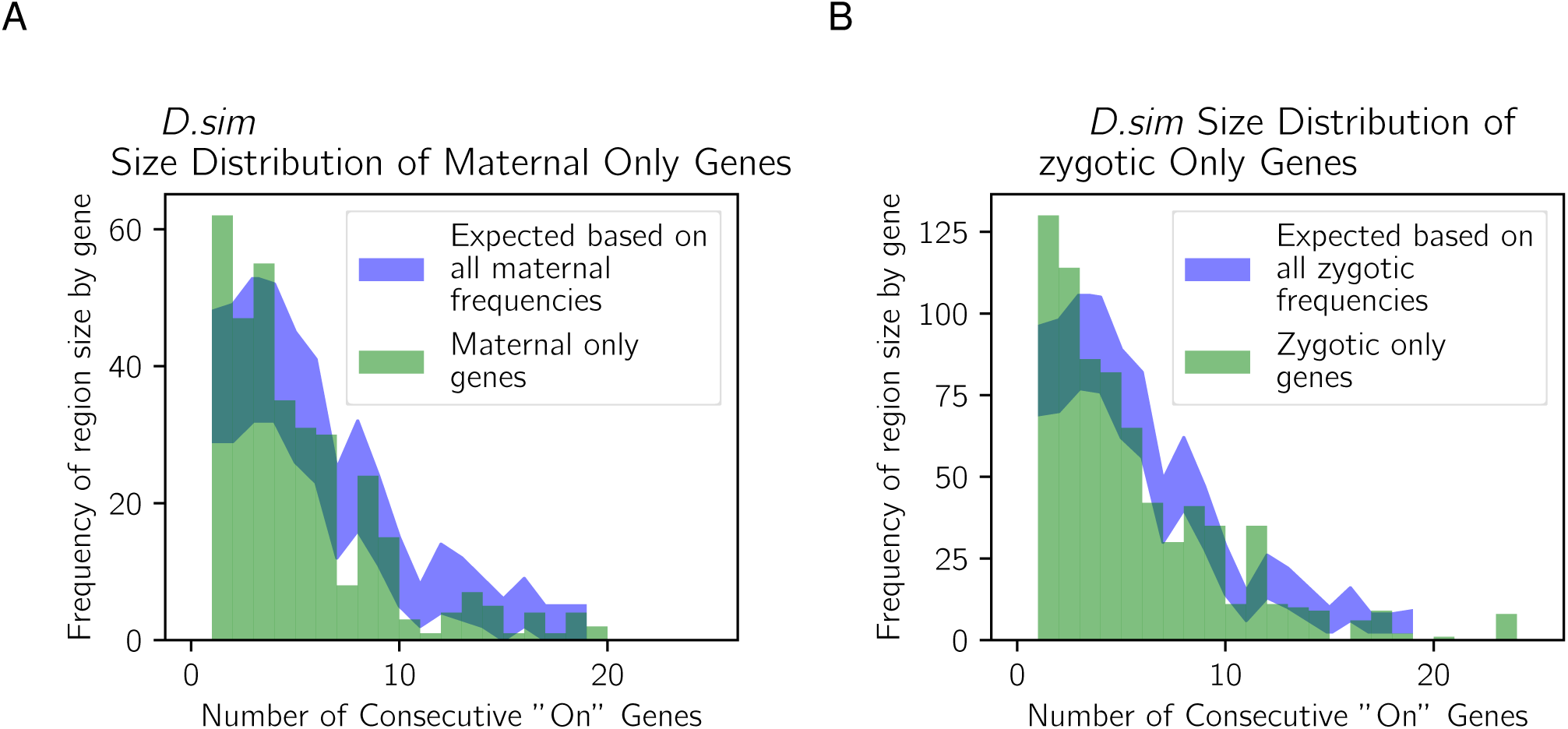
Stage-specific genes are more likely to be different from their chromatin neighborhood. *D. simulans* was chosen as a representative species. (A) Cluster size distribution of maternal-only genes (green bars) compared with the expected frequencies based on the overall cluster size frequencies observed at stage two (blue region). The expected frequencies are based on the distribution in Fig 4A multiplied by a scale factor equal to the proportion of maternally deposited genes that are maternal-only, with the shaded region representing a 95% confidence interval. (B) Cluster size distribution for zygotic-only genes (green bars) compared with the expected frequencies based on the overall cluster size frequencies observed at stage 5 (blue region) in a manner similar to Fig 5A. the shaded region represents a 95% confidence interval. For both stages, stage-specific genes are more likely to be the single gene (or one of a small number of genes) that are expressed where their neighboring genes are not, representing small numbers of “on” genes in an “off” chromatin environment.

### GC-content of upstream regions is predictive of maternal deposition

In *Drosophila*, transcription start sites are frequently associated with a spike in GC content. These spikes in GC content have been suggested to act as “genomic punctuation marks” to delineate functional regions, though their mechanisms of action are not clear[66]. To explore this phenomenon with respect to the two developmental stages we examined, we evaluated the average GC content of upstream regions for genes in stage 2 and stage 5. When comparing the GC-content of putative cis-regulatory sequences in maternally versus non-maternally deposited genes, we observed an increase in GC-content upstream of the TSS (Fig S3), as well as a dip in GC content ∼200bp upstream of these genes. In contrast, this modulation does not occur in genes that are off at both stage 2 and stage 5, nor in genes that are off at stage 2 but activated at stage 5. To determine whether this modulation of GC-content was predictive of maternal deposition, we constructed four generalized linear models using the GC-content, the motif data, and both the motif data and GC-content as data sources (see Methods). Adding the GC-content to the model that already included motif data improved the model (AIC: 185589 without GC content AIC:183079 with GC content), hence increased GC content upstream of TSS is somewhat predictive of maternal deposition, even when accounting for motif presence in this region.

The biological significance of this spike in GC content is unclear. Fluctuations in GC content have been observed in *Drosophila* previously [66], and there is evidence in humans that spikes in GC content are associated with supercoiling [67]. DNA supercoils are generated in via transcription, and positive supercoils are observed to inhibit transcription [68]. In Drosophila negative supercoils have been associated with high transcriptional activity in polytene salivary gland cells [69], and GC content directly impacts the biochemistry of DNA with respect to torsional stress [70]. As the nurse cells where maternal transcripts are produced are polyploid with a high transcription rate, nurse cell chromosomes may be under similar torsional stress. This may explain why maternally deposited genes in particular are associated with this spike in GC content.

## Discussion

Maternally deposited gene products are responsible for the first stages of embryonic development in all animals[71]. It is therefore critical that the required kind and amount of mRNAs and proteins are deposited into the unfertilized egg. Later in development, the zygotic genome becomes transcriptionally active and takes over control of development from maternal mRNAs. Failure in maternal mRNA deposition, zygotic genome activation, or the transfer of developmental control between the two genomes can lead to lethality[1,9,13], thus the gene products regulating early development are critical to organismal survival.

Previous research has shown that the maternal and zygotic mRNA expression profiles of different species of *Drosophila* are generally conserved, but with some noticeable differences[24]. To investigate the regulatory basis of transcription at these stages, we leveraged a large comparative dataset to identify the transcription factor binding motifs found in the *cis*-regulatory sequences of these genes. We found that the regulatory basis of both the maternal and zygotic-only transcripts also had significant conservation, which permitted the discovery of common features of gene regulation across *Drosophila*. Specifically, we identified transcription factor binding motifs that are associated with mRNA expression across species for maternally deposited transcripts and zygotically expressed transcripts. We also investigated the effects of other regulatory mechanisms such as chromatin state on maternal and zygotic expression of mRNAs, as well as the association of transcript levels at these two stages of embryogenesis with gene length, strandedness, and GC content.

Generally, we found a number of conserved transcription factor binding motifs associated with transcript abundance for both the maternal and zygotic-only transcripts. At the maternal stage, there were a larger number of more highly conserved motifs than were found for the zygotic-only genes. This is consistent with a previous study that found that maternal transcripts themselves were more highly conserved than transcripts at the zygotic stage [24]. Given this, surprisingly we also found less conservation of particular motifs at conserved genes transcribed at the maternal stage. As we found a number of motifs involved in regulation at the level of chromatin at the maternal stage, perhaps different combinations of chromatin regulating motifs can be utilized interchangeably without altering expression status. This could provide robustness, permitting evolutionary changes in sequence without affecting gene expression of maternal genes. In contrast, while we find that the zygotic-only transcripts are associated with fewer conserved motifs overall, and more divergent lineage and species-specific motifs, that individual conserved genes are more likely to be regulated with the same motifs. This provides conservation of gene expression by a different mechanism for the zygotic-only genes that are functionally required across *Drosophila.* Why the two stages and genomes would have such different ways of activating conserved genes across the genus is likely due to the underlying biology of regulation at the two stages, as discussed in detail below.

### Maternal Regulation

We found that motifs associated with putative *cis*-regulatory regions of maternally deposited genes are predominantly annotated as insulator binding sites. An insulator is a type of regulatory element that can block the interactions of *cis*-regulatory elements with promoters or prevent the spread of chromatin state. Insulators are known to be important in creating and maintaining the gene expression patterns, ubiquitous in *Drosophila*, and potentially a key factor for *Drosophila* to maintain such a high gene density[42]. Here, we find that the process of maternal deposition may rely heavily on insulators to express a large percentage of the genome. Because the roles and mechanisms of factors annotated as insulators are not well understood, using the term “Architectural Protein” instead of insulator binding protein may be more appropriate[72]. Recently, these proteins have been studied using genome-wide chromatin organization methods, such as Hi-C, which detects regions of interacting chromatin known as Topologically Associated Domains (TADs) and identifies boundaries between them. Histone marks appear to be enriched in certain TADs but stop abruptly at TAD boundaries, supporting the idea that certain TADs are entirely transcriptionally silenced while others are expressed[34]. Furthermore, ChIP-seq has demonstrated that TAD boundaries in other tissues are enriched in architectural protein binding sites[34], including several those that we identified in this study.

There is some disagreement on the effect that TADs have on gene expression, however. Ghavi-helm et al [73] demonstrate that the disruption of TADs does not necessarily disrupt the constituent gene expression. Instead, they suggest TAD boundaries acting to prevent interactions between TADs is rare or tissue specific. Others suggest that it is possible that TADs are increasing robustness to other regulatory mechanisms[74]. Because TAD-associated elements appear to be associated with maternal deposition in our dataset, we hypothesize that these elements are regulating maternal deposition via chromatin-level control. It is possible that there are other additional mechanisms that we do not detect.

To understand the connection between architectural proteins and maternal deposition, we need to examine where these transcripts are produced to understand the cellular context. In the ovary, nurse cells are responsible for the transcription of maternally deposited genes, and there is a considerable body of literature devoted to nurse cell biology. Much study has been directed towards elucidating how nurse cells transport their products into the oocyte and how post translational control mechanisms fine-tune protein levels of maternal transcripts[3,7,18–23,75]. However, despite this wealth of knowledge, the regulatory mechanisms by which the nurse cells specify which genes to transcribe are largely unknown. One unusual feature of nurse cells is that they are highly polyploid[76, 77]. One of the major benefits of this could be an across-the-board increase in transcription rates necessary to provision the embryo with all necessary transcripts. These transcripts represent a large proportion of the genome, with estimates ranging from 50-75%, depending on experimental conditions[3], and necessitate large amount of transcription overall in a short period of time. We extract >100ng total RNA from an embryo; this is an astonishingly large amount of RNA to be present in what is essentially at the time of fertilization a single, albeit a highly specialized, cell. One point of comparison is Abruzzi et al. 2015,[78] who extracted 2-5pg RNA per *Drosophila* neuron. A transcriptional environment that is optimized to quickly transcribe huge numbers of genes might be more amenable to control via chromatin state.

Given the amount of overlap between the motifs enriched in the *cis-*regulatory regions of maternally deposited genes and the motifs associated with TAD boundaries, it is possible that these same architectural proteins are functioning to define which genes are maternally transcribed and then deposited into the embryo. We found that the maternally deposited genes are highly clustered on the genome, which is indicative of control via architectural proteins. Additionally, we uncovered that proximity to nearby expressed genes is highly correlated with expression. We also identified a pattern whereby the relative strandedness of adjacent genes is indicative of whether they will be maternally deposited, which is a pattern that has been previously observed with insulators[34]. Each of these results is consistent with known behavior of architectural proteins, suggesting that expression at stage 2 is controlled locally on the chromosome by activating TADs rather than specific genes.

As architectural proteins are important in determining genome organization and regulating transcription to some degree in all tissues and stages, we investigated whether the regulatory patterns we observed for maternal genes were ovary-specific or shared across all stages and tissues. Many of the motif binding elements discovered in this analysis appear to be enriched in ovaries, although these proteins have important functions in other tissues as well. Some of the proteins predicted to bind our motifs have been noted for being enriched in the regulation of housekeeping genes, and as maternally deposited genes themselves are enriched in housekeeping genes, this result is perhaps unsurprising. A number of studies have suggested that in addition to the common architectural proteins shared across conditions and developmental stages, there may exist tissue-specific architectural proteins that integrate into the canonical protein complex to produce tissue-specific TAD patterns[79–81]. Perhaps this is the case with the ovary, and further study will reveal whether there are ovary-specific factors that may interact with the common architectural proteins whose binding sites we find enriched here. For example, the authors of Mataz et al. 2012[82] suggest that Shep may be a tissue-specific factor interacting with architectural proteins in the central nervous system. The enrichment Shep in the central nervous system is even less extreme than the enrichment we observe of CP190 (a known interaction partner of ZIPC, one of our maternal expression associated motifs) in ovaries, suggesting that CP190 could also qualify as tissue-specific. Alternatively, the polyploid nature of nurse cells and the extensive and rapid transcription that occurs in these cells may instead provide an extreme enrichment of the common architectural proteins, without the need for stage or tissue specific architectural proteins.

Our results show that regulation in ovaries is accomplished primarily through architectural proteins that establish general regions of open chromatin. This process can turn on a large percentage of the genome, without the need to maintain specific motifs within specific genes. However, this leaves us with the question of how the stage 2 mRNA content is so highly conserved across species overall[24], as regulation at the chromatin level would appear less precise than gene-specific regulation. Perhaps regulatory control primarily at the level of chromatin provides redundancy to maintain transcription despite the gain or loss of individual binding sites. Alternatively, there could be other levels of regulatory control that we are unable to detect, with the signal from chromatin-level control being so strong during this time. The high level of conservation of maternal transcripts is also remarkable given the importance of post-transcriptional regulators at this stage[3,19,23,83], as it is not clear if conservation at the transcript level is necessary for conservation at the protein level.

### Zygotic Regulation

Our examination of motifs that are associated with zygotic mRNA expression revealed several previously discovered motifs, including those that bind Zelda and GAGA factor (Trl). Additionally, several motifs are likely binding sites for other well-characterized developmental proteins (Table S1) which are sometimes highly localized in the embryo. If transcripts are produced in a spatially localized manner, they are necessarily not expressed in the entire embryo, and thus their signal may be more difficult to detect in our data from whole embryos. Overall, we observe few motifs at stage 5 that are conserved across species, in comparison to motifs for maternally deposited genes. However, the motifs that we do find at stage 5 tend to higher conservation within specific genes than the motifs we discover at stage 2. This highlights that it may be more important for specific genes to have precise signals after ZGA.

Additionally, in our zygotic analysis, we focused only transcripts that are present at stage 5 and do not have a maternal component, as many maternally deposited transcripts are still present at stage 5 (roughly half of maternal transcripts are still present at this stage[8,30–32]). Because many maternal transcripts are still present, analysis of the total stage 5 transcriptome would largely recapitulate the stage 2 results, especially as stage 5 transcripts are much more likely to be expressed in specific spatio-temporal patterns, which to our whole-embryo analysis would appear as low or noisy signal. Our decision to remove transcripts with maternal deposition highlights the signals that are unique to stage 5, but comes at the cost of an overall reduction in the number of genes available for analysis, resulting in higher false discovery rates for all motifs.

## Conclusions

In this study, we examined regulatory elements associated with maternal transcripts present at stage 2 of embryogenesis and zygotic transcripts present at stage 5 across species of *Drosophila*. At both stages, we found regulatory motifs that are conserved throughout the ∼50 million years of divergence represented by these species, which speaks to a conservation of regulatory mechanisms across the genus. In general, the high degree of conservation in regulatory elements at the maternal stage and the zygotic stage, while different from one another, speaks to the critical nature of the complement of transcripts present to direct early embryogenesis. The differing patterns observed in the *obscura* group species (*D. pseudoobscura* and *D. persimilis*), and the regulatory basis of changes in transcript representation between species are the subject of ongoing study. At the maternal stage, we found many regulators that appear to be defining general regions of the genome to be transcribed via chromatin regulation through architectural proteins and likely at the level of TADs. Given the exceptionally high level of conservation of maternal transcript deposition, the relatively non-specific mechanism of maternal gene regulation appears contradictory. In contrast, we found zygotic regulatory elements to be considerably more gene-specific. The different patterns of regulation for transcripts present at these two stages of embryogenesis is consistent with the specific transcriptional contexts of these two genomes, with the non-specific mechanism active in highly transcriptionally active polyploid nurse cells in oogenesis in the mother, and the gene-specific mechanism acting in the zygote where transcription is often localized in time and space.

## Methods

### Data Acquisition

RNA-seq data utilized for this study was generated previously[24], and is available at NCBI/GEO at accession number GSE112858. This dataset contains RNA-Seq data from single embryos. Embryos were collected either at stage 2, representing a time point before zygotic genome activation, and at the end of stage 5, representing a time point after widespread zygotic genome activation. Embryos were collected from 14 species, however we only used the data from 11 (*D. simulans, D. sechellia*, *D. melanogaster*, *D. yakuba*, *D. erecta*, *D. ananassae*, *D. persimilis*, *D. willistoni*, *D. mojavensis*, *D. virilis*) due to annotation deficiencies in the remaining 3. GTF files and references genomes from previously sequenced species[28] were downloaded from Flybase[84].

To determine whether a gene would be labeled as ‘off’ or as ‘on’, the overall distribution of FPKMs was analyzed. For all species, for both stage 2 and stage 5, a bimodal distribution appeared, with one peak at 0 and another at approximately e^3.5^. The commonly used cutoff of FPKM=1[85, 86] was chosen as it falls between these two distributions.

To determine which genes were orthologues, we used the FlyBase orthology table “gene_orthologs_fb_2014_06_fixed.tsv”.

### Sequence Selection

Preliminary tests were performed to determine which regions were most likely to have regulatory elements. For each gene, several regions were extracted: 10kb upstream,5kb upstream, 2kb upstream, 1kb upstream, 500bp upstream, 5’ UTR, total introns, total exons, and 3’ UTR. For each region, boundaries were obtained from the appropriate GTF and sequences were extracted using BioPython (Version 1.73,[87]). The 2kb upstream region showed the highest quality motifs (Fig S1), and thus were used for matching motifs in external databases, measuring motif overlap between species, analyzing motif position distributions, and GO analysis. For these analyses, featured in figures 1 through 3, UTRs were ignored as not every species had annotated UTRs.

### Motif Discovery

We used HOMER[29] to discover motifs in test sets using the background sets as control FASTA files, test and background sets are defined below. Deviations from the default settings include the use of the - fasta flag to specify a custom background file. For stage 2 queries, the test FASTA files included genes that had a FPKM >= 1 at stage 2 while the control FASTA files included genes that had an FPKM < 1. For the stage 5 queries, the test FASTA files contained genes where the stage 5 FPKM >= 1 and the stage 2 FPKM < 1, while the control FASTA files included genes whose stage 5 FPKM < 1 and stage 2 FPKM < 1. Additionally, we used the -p flag to utilize our computational resources more efficiently. We used - norevopp flag in the case of strand-specific searches. Motif quality was evaluated based on the HOMER-outputted q-values.

To validate the HOMER output files we used MEME[33] v4.12.0 and RSAT[88]. MEME was run using-mod zoops -nmotifs 2 -minw 8 -maxw 12 -revcomp. The RSAT analysis uses the purge-sequences tool, followed by oligo-analysis using the following parameters: -lth occ_sig 0 -uth rank 5000 -return occ,proba,rank −2str -noov -quick_if_possible -seqtype dna -l 8, followed by pattern-assembly using the following parameters: -v 1 -subst 1 -toppat 5000 −2str, followed by matrix-from-patterns using the following parameters: -v 1 -logo -min_weight 5 -flanks 2 -max_asmb_nb 10 -uth Pval 0.00025 -bginput - markov 0 -o purged_result.

### Stage-specific gene analysis

For analyses of zygotic transcripts, such as the motif analysis, we defined genes as being zygotic-only if they were off at stage 2 (FPKM <1) and on at stage 5 (FPKM >1), for N=10,215 genes across all species. It is necessary to impose such a restriction, as a large percentage (approximately 85%) of genes that are zygotically expressed were also maternally deposited, and analysis of stage 5 regulatory mechanisms would be confounded the signal of stage 2 genes. For analyses of maternal-only transcripts, we define maternal only if they are on at stage 2 (FPKM >1) and off at stage 5 (FPKM <1). As the class of maternal-only genes is very small (N=3194 across all species), we were unable to obtain results for some analyses such as the motif content detection and GO analyses for this group of genes.

### Motif Sharing

To determine weather motifs were shared between species, the HOMER-formatted motifs were converted to meme-formatted motifs using chem2meme from the MEME Suit[33]. Tomtom, also from the MEME Suit, was then used to find matching motifs, using default parameters. For a motif to be considered shared with another species, the Tomtom output threshold of *α* = .05 was used. this technique was used to calculate the similarity of motifs found in different species, as well as to evaluate the similarity of different motif discovery strategies using MEME, RSAT, or HOMER with alternative parameters.

To refine the results of shared motifs, we applied an additional quality cutoff. For stage 2, motifs were first filtered for a q-value of less than 1e-100, and for stage 5, motifs were first filtered for a q-value of 1e-10. The difference in the cutoffs used at the two different stages was due to the differences in the overall distribution of q-values for these stages due to a reduced number of zygotic-only genes (see zygotic-only motifs above).

Because sharing was calculated on a by-species basis, it is possible that one species has a motif that meets the criteria for being shared among all other species while other species’ version of that same motif failing to meet the criteria. This can occur, for example, when a motif is an intermediary version of two motifs that fall just outside the cutoff.

To find proteins that bind to the discovered motifs, we used Tomtom to query JASPAR and Combined Drosophila Databases using the default parameters[89].

### Motif Position and Count

Motif position was determined by using the scanMotifGenomeWide tool to in the HOMER package. Queries were performed by scanning the discovered motifs against the fasta files for each gene. The 5’ boundary of the motif was used as the motif position. For the motif counts per gene used in many downstream analyses analysing motif position distributions, GO analysis, GC content analysis, and motif strand analysis. We used this output and counted the occurrence of a given motif in the target region. To quantify positional distribution similarity, we used the stats.anderson_ksamp function from the scipy library V1.2.1[90]. Distributions were considered to be different at *α* = .05 after Bonferroni correction.

### Transcript Enrichment by Tissue

Expression data for various adult tissues was downloaded from modENCODE[57]. To compare enrichment for transcripts with different magnitudes of abundance, we applied an additional normalization. For each transcript, transcript levels in FPKMs were divided by a scaling factor equal to the average of the expression levels in ovaries. This normalization preserves the relative abundances within each transcript, but allows for visualization of transcript levels with dramatically different overall expression levels.

### Housekeeping Gene Identification

To compare the enrichment of the discovered motifs in maternally deposited genes versus housekeeping genes, we identified housekeeping genes using modENCODE data [57]. Housekeeping genes were defined as having expression in each of the following tissue types: larval CNS, larval ovaries, male larval gonads, male reproductive tracts, adult midguts, adult heads. In addition, putative housekeeping genes needed expression levels of greater than 1 FPKM in our stage 2 and stage 5 dataset in *Drosophila melanogaster*.

### Expression by Position

*D. melanogaster* expression data by position was downloaded from modENCODE[57] for several tissue types. Positions for each motif was determined as previously described in the Motif Position and Count section above. For each instance of the motif of interest, we determined expression values in area from −10kb to +10kb. Transcript abundance in FPKMs were then normalized by the average FPKM reported on the track.

### GO Analysis

We used the R package clusterProfiler 3.10.1[51] and the org.Dm.eg.db 3.7.0[91] dictionary to perform gene ontology (GO) analysis. For the stage 2 comparison, we generated a test set of the melanogaster gene names for every gene in our dataset that was maternally deposited in at least any 7 of our species, and performed an enrichment analysis using enrichGO’s default parameters using a background set of all *D. melanogaster* genes. For the stage 5 comparison, we generated a test set of the *D. melanogaster* gene names for which at least two orthologues in our dataset showed zygotic-only expression (see Zygotic-only motifs section above for definition). This threshold approximates the percent of the genome that we observed to be zygotic-only. We then performed an enrichment analysis using enrichGO’s default parameters using a background set of *D. melanogaster* genes that are not maternally deposited in at least two species. This analysis therefore specifically examines the zygotically activated genes in the context of genes that are “off” at stage 2 (FPKM<1 at this stage). For our analysis of stage 2 motifs, we generated a test set for each motif consisting of genes that contained that motif in at least two species and were maternally deposited (FPKM > 1) in at least two species. We then performed an enrichment analysis using enrichGO’s default parameters using a background set of all *D. melanogaster* genes. For our analysis of stage 5 motifs, we generated a test set for each motif using genes that were represented by transcripts >1 FPKM at stage 5 in at least two species and had the motif of interest in at least two species. We then performed an enrichment analysis using enrichGO’s default parameters using a background set of *D. melanogaster* genes that were represented by transcripts >1 FPKM at stage 5. To visualize our results, we employed the dotplot method for enrichGO objects, also from the clusterProfiler package. For each motif, the top 3 GO terms were identified and added to the y-axis labels. Whenever any GO category from another motif was identified as statistically significant (*α* = .05), that GO category was shaded appropriately.

To discover motifs associated with particular GO categories, we generated a list of genes that were both maternally deposited and associated with each GO term of interest, as well as a list of genes that were maternally deposited but not associated with the GO term of interest. For each GO term, we ran HOMER using the same parameters as the initial motif discovery, using the genes associated with the GO term as the test list and the genes not associated with the GO term as the background. We restricted this analysis to the upstream regions of *Drosophila melanogaster* genes.

### Model Fitting

Logistic regression was performed using the “glm” function in R, using the logit link function. As inputs, we used the list of motifs generated from HOMER and their counts as described in the “Motif Position and Count” section above. To avoid redundant motifs in our model, only motifs of size 10 were considered. To evaluate the strand-specificity of motifs, we compared two generalized linear models using the formulas indicated in S1_Model_Generation.pdf. To identify the most important motifs, the R function stepAIC from the MASS library 7.3-51.4[92] was used to find generate an ordered list of motifs. The base model used contained no additional features (chromatin state, etc). StepAIC was run 8 steps to generate a short list of motifs for evaluation.

### Analysis of physical clustering of co-expressed genes

To evaluate the effect of gene cluster size on expression, we iterated through each species for both stage 2 and stage 5 and assigned sizes of co-expressed gene clusters on the chromosome, based on how many adjacent genes were coexpressed, resulting in cluster size frequencies for each genome. Errors were calculated using 95% confidence interval for a two-tailed binomial distribution.

To compare the clustering of different datasets with varying percents of “on” genes, we employed the Wald–Wolfowitz runs test.

### Tissue-specific RNA Levels

modENCODE tissue profiles[53] were downloaded from flybase.org. Flyatlas2 tissue profiles were downloaded from http://flyatlas.gla.ac.uk/FlyAtlas2/[54].

### Gene length

To determine gene length, we examined the relevant line of the appropriate .GFF file and took the difference between the end and the start positions.

### Distance between genes

To determine the distance between genes, we look at the appropriate .GFF file and took the difference of positions between adjacent genes from transcription start site (TSS) to TSS.

#### Maternal deposition rates as compared to gene length, distance, and orientation

Genes were binned by category and by either distance or length. For the top plot, 150 bins of 70bp width were used. For the bottom plot, 60 bins of 70bp width were used and bins with fewer than 6 genes were disregarded. confidence intervals were calculated using the binomial distribution with *α* = .05 after Bonferroni.

### GC content

GC content levels associated with each gene were evaluated by calculating the number of GC nucleotides within a sliding window of size 50bp for each of 1950 window positions to cover the upstream 2kb of each gene. To evaluate the first bin of each gene, the region from −1bp to −50bp was extracted, and the number of G and C nucleotides was counted. The result was divided by 50 to get the %GC for this window. To calculate the GC content for the next bin, this process was repeated on the region from −2bp to −51bp. Each bin had its GC content evaluated this way until the final bin of −451bp to −500bp. To evaluate how closely a particular upstream region resembled a maternally deposited-like distribution or a non maternally deposited-like distribution for the purposes of modeling, we calculated the average GC content for each position of maternally deposited, and not maternally deposited genes. Then for each gene, we measured the correlation between the GC content and that of both category averages. We used the difference in these correlations as a metric to evaluate similarity in GC content for each gene.

## Acknowledgements

We would like to thank Joel Atallah for his work on the original dataset, Gizem Kalay, Anna Feitzinger, and Emily Cartwright for comments on the manuscript, and all members of the Lott Lab for feedback.

## Supporting Information Figure Captions

**S1 Table:** A summary of the top ranked zygotic motifs. Motifs were selected if had enrichment if they were enriched in the combined upstream regions of all species with a q-value < 1e-50 and a Tomtom match to any motif in an existing database with q < .1. If there were more than one, the best two matches to motifs in existing databases were reported in the Best Match column. Some motifs are plausible binding sites for known embryonic regulators.

**S1 Fig:** Distribution of motif qualities by location in a representative species in each stage. *D. ananassae* was selected as a representative species. Motif qualities are given by the negative natural logarithm of the q-value outputted by HOMER. High quality motifs enriched for stage 2 (A) are most likely to be found in the 2kb upstream of a gene. Motifs for stage 5 are generally less high quality by this metric, and while the highest quality tend to also be enriched 2kb upstream, some are enriched in 2kb upstream regions of non-expressed genes or enriched in exons.

**S2 Fig:** Representative positional distributions of motifs. Distributions for both maternally deposited genes (“on”) and non-maternally deposited genes (“off”) are shown. (A) The positional distribution of the DREF motif, which follows the same pattern as M1BP, Zipic, Ohler-6, and E-box, and many motifs without identified factors that bind them. These motifs are found upstream of maternally deposited genes (red), with a higher frequency closer to the transcription start site. They are not found with any frequency in non-maternally deposited genes (blue). (B,C) Positional distribution patterns of some rare, undocumented motifs. In both, we see that the motif is more enriched in maternally deposited genes than in non-maternally deposited genes, but that the enrichment difference is less than those motifs represented by (A) above. In (B), this motif is most highly enriched upstream, less enriched around the transcription start site (TSS), and more highly enriched again downstream of the TSS (though less so than upstream). In (C), we see the highest enrichment downstream of the TSS, with a dip in enrichment around the TSS, and less enrichment upstream of the TSS than downstream

**S3 Fig:** GC content of the region upstream of the TSS. GC content for each gene in a sliding window with 50bp width is summed for each gene in the category. (A) Maternally deposited genes. (B) Non-maternally deposited genes. (C) Zygotic-only genes. Note the high number of genes with higher GC content immediately upstream of maternally deposited genes, and the lower GC content upstream of this GC-enriched region.

S4 Fig: Low quality motifs are less likely to be shared across species. In a manner similar to figure 1 a and b, we discovered motifs for each species at both stage 2 and stage 5 and evaluated what percent of motifs were shared among species. Unlike the analysis described in figure 1 a and b, we did not apply a quality filter.

S5 Fig: GO-term specific motifs exist, but are not predictive of maternal deposition. The effect and p-value column data are generated from a generalized linear models of the form [maternal deposition] ∼ [motif presence], given a number of genes whose adjacent genes are expressed. Although the effect is always positive, indicating a slight increase in maternal deposition rates for genes with this motif, the high p-values indicate that these results are not statistically significant.

S6 Fig: maternal deposition is a more important attribute for these genes than housekeeping. (A) within non-housekeeping genes, the discovered motifs are much more common within maternally deposited genes. Error bars represent 95% confidence intervals by the binomial distribution. P-values are generated by the prop.test function in R. (B) genes labeled as maternally deposited are more likely to contain these motifs than genes labeled as housekeeping. effects were calculated by generating a generalized linear model in the form [presence of motif within genes] ∼ [housekeeping or not] + [maternally deposited or not]. Error bars represent standard error.

