## Supplementary material for "The conserved regulatory basis of mRNA contributions to the early *Drosophila* embryo differs between the maternal and zygotic genomes": S1 File

**Model Generation**

To generate the models used evaluating the effect of motif strand, we generated a dataframe with rows corresponding to each gene, and columns containing whether or not it was maternally deposited named "bool_stage2", each stranded motif, and each unstranded motifs. Additionally, we created two lists containing the names for these motifs, named "unstranded_motifs" and "stranded_motifs" respectively. The contents of these lists are available in "S2_unstranded_motifs.txt" and "S3_stranded_motifs.pdf", respectively. We then constructed the model using the following code to generate the formula used in the glm:

```
unstranded_motifs_list <- paste(unstranded_motifs,collapse = "+")
unstranded_formula <- as.formula(paste("bool_stage2~+", unstranded_motifs_list))
stranded_motifs_list <- paste(stranded_motifs,collapse = "+")
stranded_formula <- as.formula(paste("bool_stage2~+",stranded_motifs_list))
```

Similarly, to evaluate the benefit of including GC content into the model, we generated two models using the following formulas:

```
motif_list <- paste(stage_2_motifs, collapse = "+")
just_motifs_formula as.formula(paste("bool_stage2~",motif_list))
both_formula <- as.formula(paste("bool_stage2~","GC_score +",s2_apart))
```

Where "stage_2_motifs" is the list of motifs considered in the model, available in "S4_GC_motifs.txt" and "GC_score" is the metric used to measure how closely a given upstream region's GC content resembles the average GC content of a maternally deposited gene verses a non maternally deposited gene.
